# rRNA Expansion Segments Mediate Ribosome Dimerization as a Conserved Stress Response

**DOI:** 10.64898/2026.02.21.707218

**Authors:** Wenhong Jiang, Chen Chen, Xing Wang, Wei Huang, Dawid Krokowski, Ziyao Chen, Jiahao Xie, Zhaoming Su, Maria Hatzoglou, Derek J Taylor, Qiang Guo

## Abstract

Inhibition of mRNA translation is a common feature in proteostatic stress cellular responses. Puromycin, a widely used compound for studying translation, disrupts protein synthesis by mimicking the 3’ end of aminoacyl-tRNAs. Despite its extensive use as a research tool to probe the connection between translation activity and various physiological and pathological states, the cellular response associated with puromycin-induced translation stress remains incompletely understood. Utilizing in situ cellular electron tomography and topology analysis, we visualized the translation machinery at high resolution. Our analysis revealed that puromycin-treated neuronal cells exhibit an accumulation of "idle ribosomes" characterized by the binding of eIF5A, indicating a close association of this factor with translationally inactive ribosome states under stress. Additionally, the idle ribosomes formed dimeric complexes mediated by ribosomal RNA expansion segments, suggesting an evolved mechanism involving these regions in translation hibernating and protecting idle ribosomes. We show that the hibernating disome formation is not unique to puromycin administration but represents a conserved mechanism as a response to different cellular stressors including those associated with endoplasmic reticulum (ER) stress and amino acid depletion. Altogether, our findings shed light on previously unexplored aspects regarding unique states of mammalian ribosome hibernation, and collectively offers new avenues for understanding the correlation of cellular stress response and the regulation of protein synthesis.

## Introduction

Ribosomes are the protein synthesis machines in all forms of life, translating messenger RNA (mRNA) into proteins ^1^. During this crucial process, ribosomal complexes undergo several distinct stages that include initiation, elongation, termination, and recycling. The detailed mechanisms of the translation process, particularly those steps involving elongation factors, have been elucidated primarily *in vitro* ^2–5^. Recently, advancements in cryo-electron tomography (cryo-ET) have enabled the direct visualization of the translation landscape in its physiological setting. This cutting-edge technique facilitates the determination of high-resolution ribosome structures under both homeostatic conditions and diverse stress stimuli, providing unprecedented insights into translation regulation mechanisms within the context of living cells ^6–10^.

Unlike conventional ribosome-targeting antibiotics that directly inhibit translation elongation, puromycin mimics the 3’ end of aminoacyl-tRNA. This molecular mimicry allows puromycin to be incorporated into the ribosomal A-site during elongation, where it terminates protein synthesis through premature covalent binding to nascent polypeptide chains ^11,12^. While this property disqualifies it for therapeutic application, the unique puromycylation reaction facilitates the development of puromycin-based techniques including quantitative assessment of translation kinetics ^13^, spatiotemporal mapping of translating ribosome distributions ^14^, and affinity-based isolation of nascent polypeptide ^15^, providing critical insights into translation regulation and proteome dynamics. Despite its widespread use, however, the precise effects of puromycin on the translation processes and its cursory roles including effect on cellular response remain a subject of ongoing investigation, sparking debate about the fidelity of established puromycin-based techniques ^16–18^.

Using primary cultured neuronal cells, we employed cryo-ET and subtomogram averaging to visualize the translation landscape following puromycin treatment. This approach enables the quantitative characterization of translation machineries and their topology under translational inhibition, including the identification of a non-translating ribosome complex composed of eIF5A, SERBP1, and eEF2. We also identified topologies of ribosome dimers, or disomes, where two ribosomes are tethered together by kissing stem-loops coordinated by ribosomal RNA (rRNA) expansion segments. These disomes are distinct from those reported in bacteria ^19,20^ and eukaryotic parasites ^21^ that involve defined hibernating factors. We further show that these hibernating, dimeric ribosomes similarly and reversibly form in response to endoplasmic reticulum (ER) stress inducers, including cyclopiazonic acid (CPA) administration and amino acid deprivation. These findings enhance our understanding of the intricate mechanisms regulating translation *in situ* and offer new insights into cellular responses to translational stress.

## Results

### Puromycin reshapes the translation landscape in neuronal cells

To unravel the translation landscape *in situ*, we employed cryo-ET on primary cultures of rat hippocampal neuron cells. Tilt series were acquired from lamellae prepared via cryo-focused ion beam (FIB) milling in the soma region of the cells (Figure 1A-B, Figure S1). To evaluate the effects of puromycin on the neuronal translation landscape, we analyzed both control neurons (161 tomograms) and neurons treated with puromycin for 10 minutes (164 tomograms). Such treatment duration did not induce any discernible cell death, but significantly reduced translation activity, as confirmed by significant reduction in polysomes and an increase in free 80S ribosomes and an erupted disome peak (Figure S1F).

**Figure 1.**
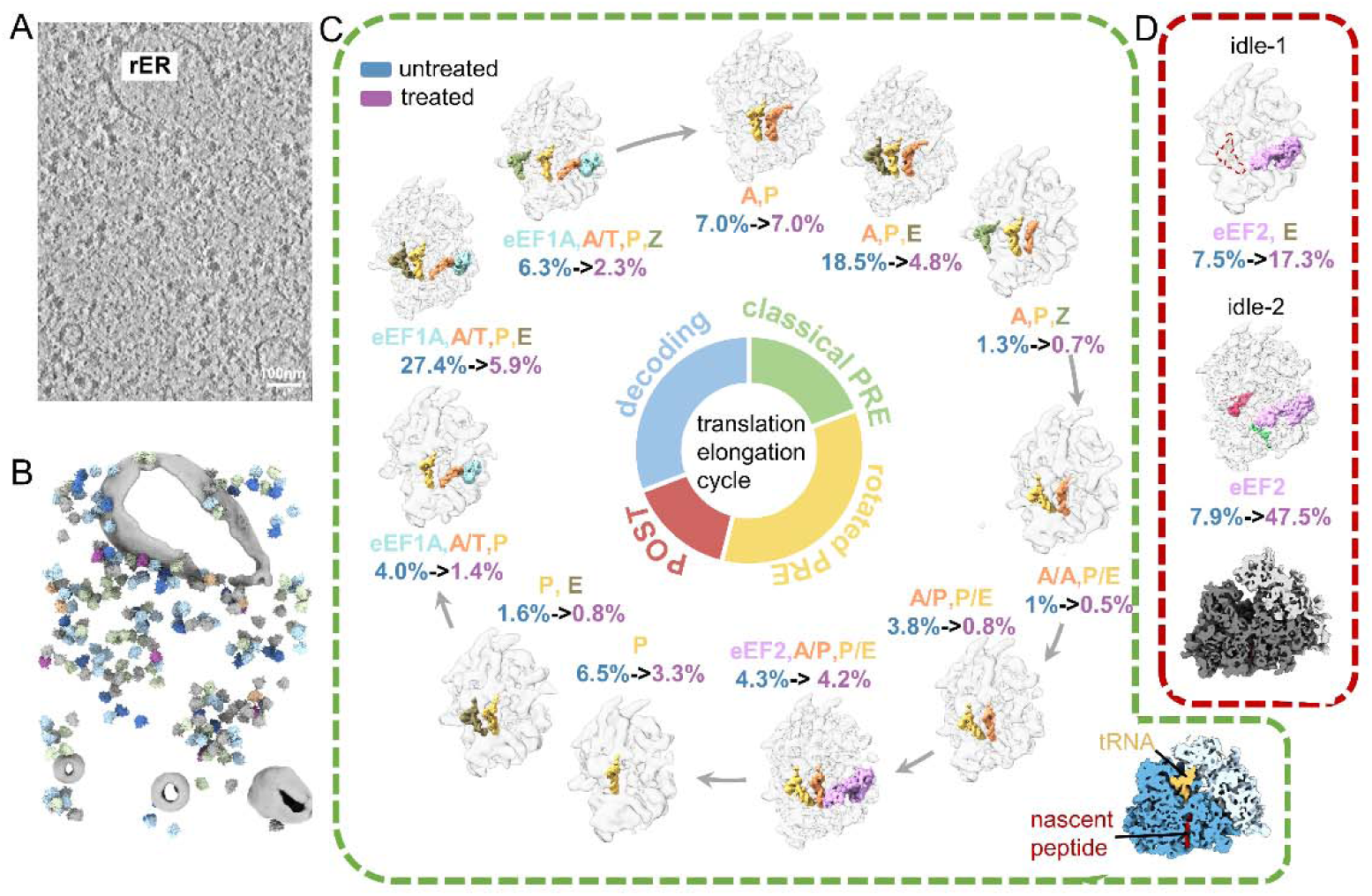
Puromycin reshapes the translation landscape. (A) A cross-section of a representative tomogram showing the cytosolic region of a neuron cell. Scale bar:100 nm. (B) The segmentation of (A) with ER colored in grey. Ribosomes were pasted using the position and orientation information from subtomogram averaging. The ribosomes were color-coded according to translation states using the color scheme corresponds to Figure3B. (C) The inferred translation elongation cycle of translationally active ribosomes. Percentage values correlate with ribosome abundance (blue: untreated group, purple: puromycin-treated group). Bottom: cross-sectional view of direct reconstruction of all the translationally active ribosomes with P-tRNA in yellow, nascent peptide in red, large subunit in dark blue and small subunit in light blue. (D) Two conformations of translationally inactivate ribosomes. For idle-1 state, the ribosome bound with E-tRNA and eEF2 (pink) exhibits reduced density for the E-tRNA, as indicated by a dashed contour in the map. For idle-2 state, E-tRNA (grey) and associated ribosomal factors (pink, green, red) are shown. Bottom: cross-sectional view of direct reconstruction of all translationally inactivate ribosomes (the large subunit: deep grey, the small subunit: light grey).

Ribosomes and subunits were identified within the reconstructed tomograms. Extracted ribosome particles underwent rigorous classification and averaging procedures (Figure S2). A total of 30,071 80S ribosomes were isolated and subjected to subtomogram averaging to yield map of the 80S ribosome at an overall resolution of 4.8 Å (Figure S1D-E), confirming high data quality.

The functional states of 80S ribosomes were classified through integrative analysis of inter-subunit rotation and occupancy of the three canonical (A-, P-, E-) tRNA binding sites (Figure 1, Figure S2). Thirteen distinct ribosomal states were defined and individual structures solved at resolutions ranging from 6.8 Å to 17.9 Å. Among these, 11 structures exhibiting canonical P-site tRNA engagement combined with variable A/E-site configurations (Figure S3A). Continuous density extending from P-site tRNA through the peptide exit tunnel was observed in these classes, consistent with nascent peptide occupancy during active elongation (Figure 1C). These states correspond to established intermediates of the translation elongation cycle ^2,7,9^, including a predominant decoding state, classical and rotated pre-translocation (PRE) states, and the low-abundance post-translocation (POST) state. Notably, ribosomes containing Z-site tRNA showed equivalent upstream and downstream neighboring densities, contrasting with previously reported downstream ribosome clustering in mouse embryo fibroblast cells ^7^. This discrepancy challenges the notion that Z-site tRNA is a feature of translational stalling.

Additionally, two distinct states were identified. Each of these states lacked P-site tRNA and nascent peptide density but retained eEF2-bound in the ribosomal A-site (Figure 1D); both of these features are characteristic of previously described idle ribosomes ^22^. Supporting the assignment, neighboring ribosomes were detected on both mRNA entry and exit sides of the 11 elongation-associated states, whereas the two idle states exhibited either single or no neighboring densities (Figure S3B).

To visualize puromycin’s effects on the translation landscape, these classified ribosome states were further separated based on treatment. As anticipated, 10-minute puromycin treatment significantly reduced actively elongating ribosomes from 81.7% to 31.7%, concomitant with an approximately fourfold increase in idle ribosomes (15.4% → 64.8%) (Figure 1C-D). Unlike other antibiotics that directly block the elongation step of translation, puromycin’s aminoacyl-tRNA mimicry induces premature termination ^11^. While this termination-specific action results in an increased population of idle ribosomes, no substantial changes in elongation-associated ribosome states were detected under these conditions (Figure S4). Furthermore, puromycin exhibited minimal effects in separating ribosomes into small (5.6% treated vs. 5.5% untreated) and large (7.1% treated vs. 5.5% untreated) subunits, suggesting that it neither promotes ribosome dissociation nor markedly alters the relative abundance of initiation-associated ribosome populations. This observation aligns with finding in prokaryotes where puromycin-mediated peptide release preserves ribosome monomer integrity ^12^, contrasting with antibiotics that directly impede translation elongation ^6,7^.

### Puromycin induces idle ribosomes by recruiting eIF5A

Our cryo-ET analysis resolved two distinct idle ribosome states accumulated after puromycin treatment (Figure 1D). However, the resolution was insufficient to unambiguously identify features in the map to suggest additional factors were bound to the idle ribosomes. Thus, to investigate potential contributing factors, we applied cryo-EM to the cellular lamellae using GisSPA ^23^ to generate cryo-EM maps with superior resolution and enhanced structural features (Figure S5).

The first state (idle-1), was characterized by the occupancy of eEF2 and E-tRNA (Figure 2A), and is comparable to a previously documented state of translationally inactive ribosomes ^24^. The second state (idle-2), comprising nearly half of the ribosome population after puromycin treatment, reveals a novel assembly composed of eEF2, SERBP1, and eIF5A (Figure 2B). Interpreting the fitted structure highlights the positioning of eIF5A to span the ribosomal E-P sites and exhibits well resolved features in the map near the peptidyl transferase center. Additionally, eIF5A contacts ribosomal protein uL1, inducing an inward movement of the L1 stalk toward the central protuberance (Movie S1).

**Figure 2.**
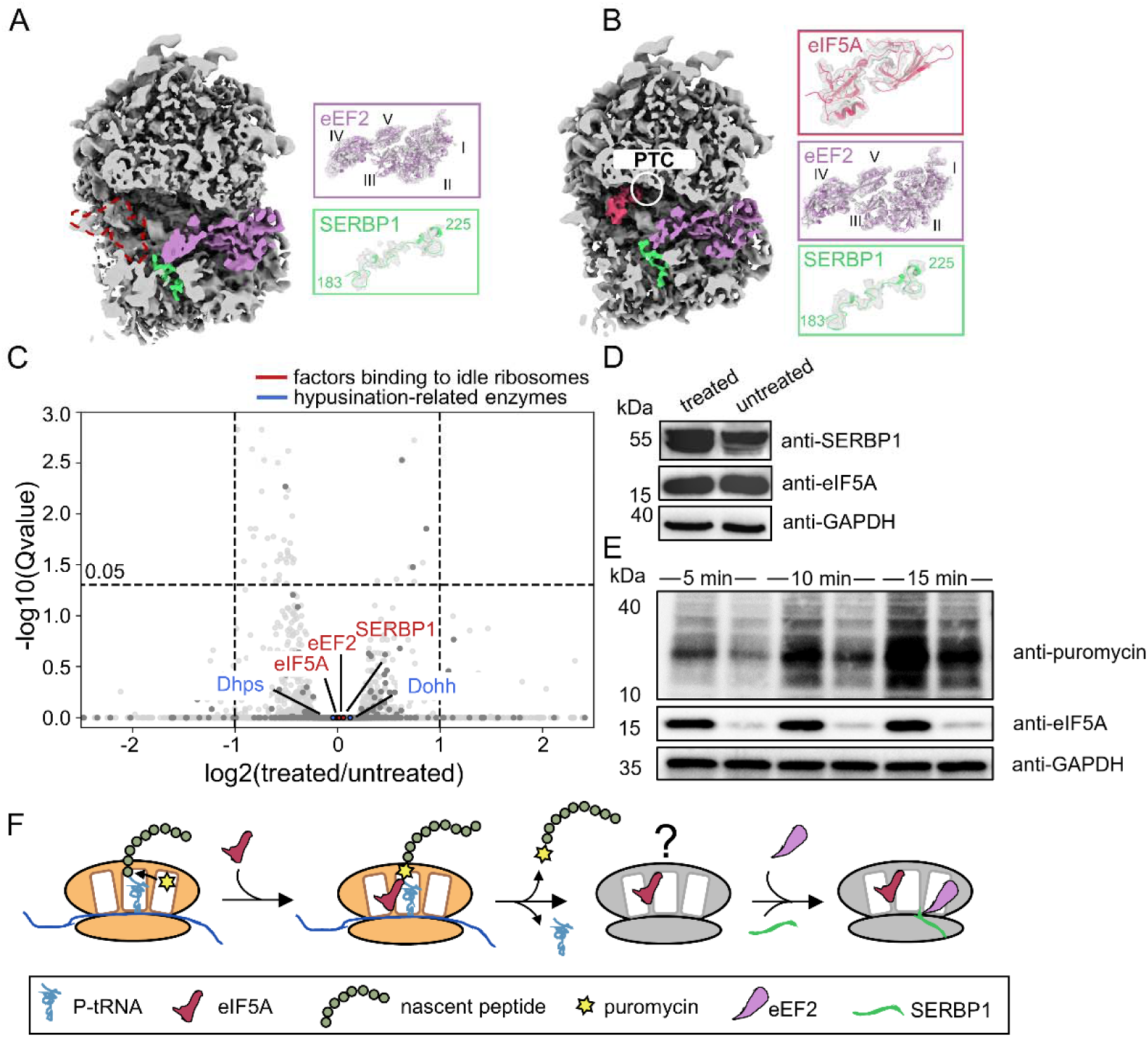
Idle ribosomes were enriched after puromycin treatment. (A) A cross-section of the idle ribosome density map (idle-1) with eEF2 in purple and SERBP1 in green. The ribosome exhibits reduced density for the E-tRNA, as indicated by a dashed contour in the map. (B) A cross-section of the idle ribosome density map (idle-2) with eIF5A in red, eEF2 in purple and SERBP1 in green. The terminal end of eIF5A is positioned within peptidyl transferase center (PTC) region. (C) The effect of puromycin on transcription. The factors shown in B) are highlighted with red and hyusine modification-related enzymes of eIF5A are with blue in the volcano plot. (D) Western blot analysis of factors shown in B). (E) Western blot detection of puromycylated nascent peptides with/without eIF5A knockdown. (F) A proposed model delineating eIF5A’s involvement in idle ribosomes formation.

Puromycin assays conducted *in vitro* initially identified eIF5A as a translation initiation factor that participates in the formation of the first peptide bond ^25^. Like its bacterial homolog EFP, eIF5A plays a crucial role in translating slow polyproline sequences, which act as poor peptide donors during translation, by positioning the CCA-end of the P-tRNA for transfer of the nascent chain to the aa-tRNA ^26–28^. Recent studies in oocytes have demonstrated that eIF5A is recruited to dormant ribosomes by Dap1, alongside eEF2 and Habp4, a paralogue of SERBP1. This recruitment highlights eIF5A’s diverse roles beyond translation initiation and elongation, including its involvement in managing inactive ribosomes ^29^. In our structure, SERBP1 occupies the mRNA entry channel and is proximal to bound eEF2. Thus, together, eEF2 and SERBP1 form a conserved spatial configuration that is observed in dormant ribosomes across diverse species ^6,30^.

To further interrogate eIF5A binding to idle ribosomes, we conducted RNA-seq and Western blotting (Figure 2C-D) experiments. These analyses revealed that eIF5A and its regulatory enzymes remain unchanged upon puromycin treatment, suggesting that eIF5A is actively recruited rather than being a consequence of increased expression. As puromycin is a poor peptide acceptor ^31^, one possibility is that eIF5A recruitment reflects principles similar to those described for ribosomes encountering inefficient peptide acceptors. The release of puromycin-modified peptides is consistent with the accumulation of translationally inactive ribosomes retaining eIF5A binding and lacking detectable P-site tRNA density. Consistent with this hypothesis, we observed that eIF5A knockdown reduces the incorporation of puromycin into newly synthesized peptides (Figure 2E), a finding consistent with observations in other genetic knockout models ^32^. Together, our results support a model in which slowed elongation by the ribosome, due to poor peptide acceptors, is coincident with increased recruitment of eIF5A. In conjunction with SERBP1 and eEF2, eIF5A induces idle ribosome accumulation by blocking all functional sites (A-, P-, E-sites and the mRNA channel) of the translation machinery (Figure 2F).

### Puromycin induces ribosome topological distribution alternation

The *in situ* tomographic data enables the investigation of translation regulation at the level of ribosome spatial distribution or topology rather than individual translation machineries. Therefore, we performed neighboring ribosome topology analysis using the NEMO-TOC method we developed ^33^. Nearest-neighbor distance analysis revealed no significant differences before and after puromycin treatment, consistent with the finding that the release of puromycylated peptides does not split 80S ribosomes (Figure S6).

The spatial organization of neighboring ribosomes was further clustered into five representative clusters, encompassing 56% of all 80S ribosomes (Figure 3A). Two of these clusters, which could be extended to form helical and planar superstructures, represent the basic units of cytosolic and ER-bound polysomes, respectively. The remaining three clusters exhibited features of unextendible ribosome dimers due to steric clashes and were named Disome 1, Disome 2, and Disome 3. Among these, the Disome 1 interface is mediated by the body of the small subunit, as previously documented ^33^. The other two clusters represent novel dimers mediated by large subunits, which will be discussed in detail later.

**Figure 3.**
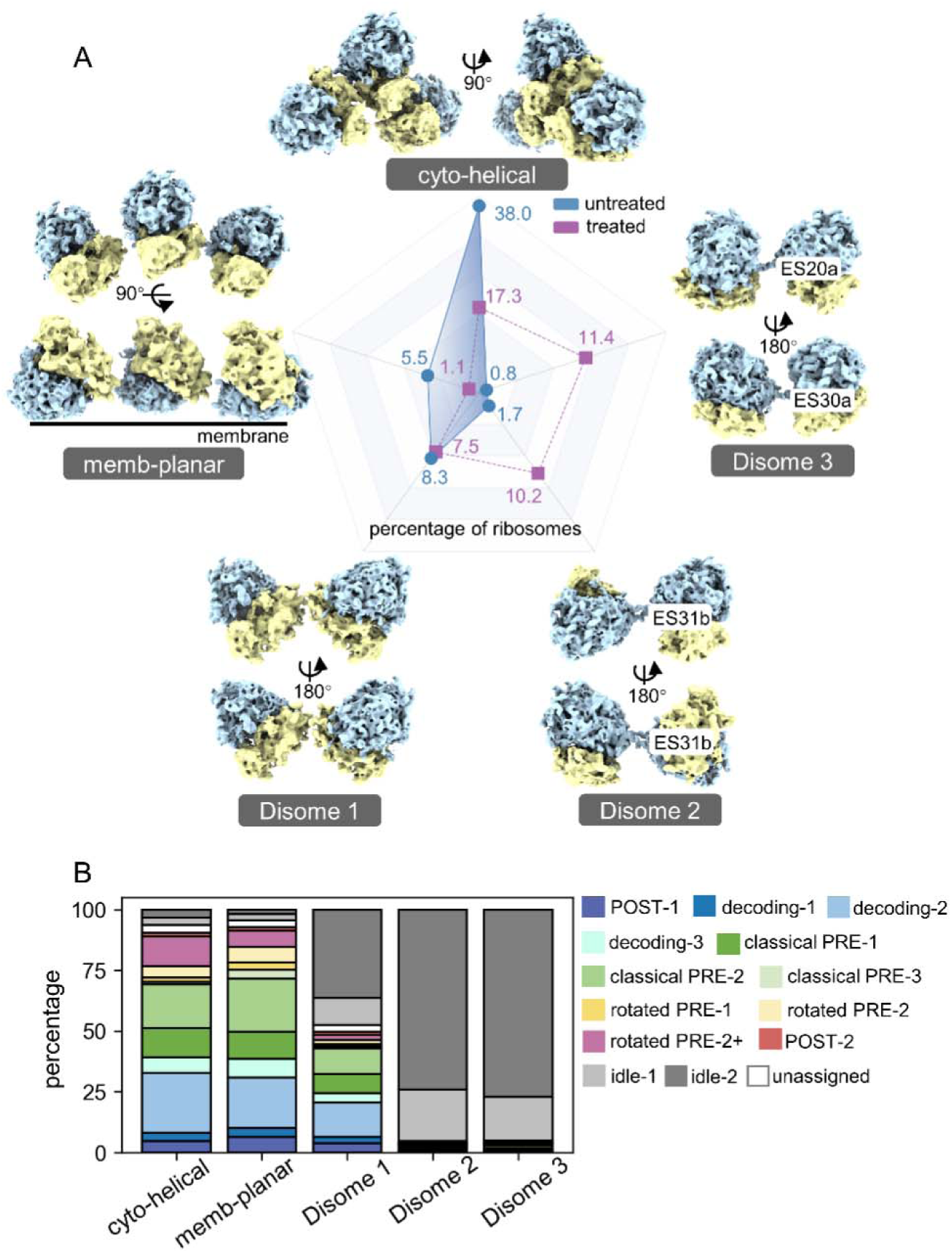
Topological alternation of neighboring ribosomes upon puromycin treatment. (A) Five representative topologies of neighboring ribosomes with corresponding abundance distribution before/after puromycin treatment. (B) Translation states distribution of ribosomes in different topology groups.

We subsequently analyzed whether ribosome spatial organization correlates with overall translation activity. Indeed, puromycin treatment exhibited differing impacts on the populations of these five clusters. While the two typical polysome configurations were significantly repressed in all populations, levels of the two dimers mediated by large subunit interactions, Disome 2 and Disome 3, increased from less than 2% to more than 10% upon puromycin treatment (Figure 3A). This distinct pattern of topological cluster alternation underscores a strong coupling between translation activity and ribosome neighborhood spatial organization. Specifically, when examining individual ribosome states within each neighboring cluster, we observed that ribosomes in cyto-helical and membrane-planar clusters were predominantly engaged in elongation cycles, consistent with previously characterized polysomes (Figure 3B). Conversely, the two large subunit-mediated dimers, the Disome 2 and Disome 3, were exclusively composed of idle ribosomes, indicating they represent higher-order topologies for translationally inactive ribosomes. In contrast, dimers mediated by small subunit interactions (i.e. Disome 1) displayed no preference regarding ribosome translational activity, aligning with their unaltered population upon puromycin treatment (Figure 3B).

### Novel hibernating disomes are maintained by rRNA expansion segment interactions

Our observation of translationally inactive ribosome dimers, Disome 2 and Disome 3 echoes previous descriptions of collided-stalled or hibernating disomes ^34^. The unoccupied P-site, alongside the spatial arrangement of these neighboring ribosomes argues against the collision hypothesis, due to the substantial distance separating their mRNA tunnels (Figure S7A, B). Consequently, the disomes observed herein more likely represent a novel mechanism of ribosome hibernation. In contrast to previously characterized hibernating dimers, which are mediated by SSU-bound hibernating factors ^19,20^, ribosomal proteins of SSU and LSU ^35^, or SSU itself ^21^, hibernating disomes observed in this study are mediated by eukaryotic specific rRNA expansion segments of the LSU.

Although the overall resolution of our disome structure is somewhat limited due to inherent flexibility between the two ribosomes, focused refinement enables improved resolution to facilitate fitting atomic models of individual ribosomes for further analysis. Our data reveal that disomes exhibit sequence-specific inter-ribosome interactions, coordinated by a pseudoknot formed through base-pairing between expansion segments of neighboring 28S rRNA molecules (Figure 4).

**Figure 4.**
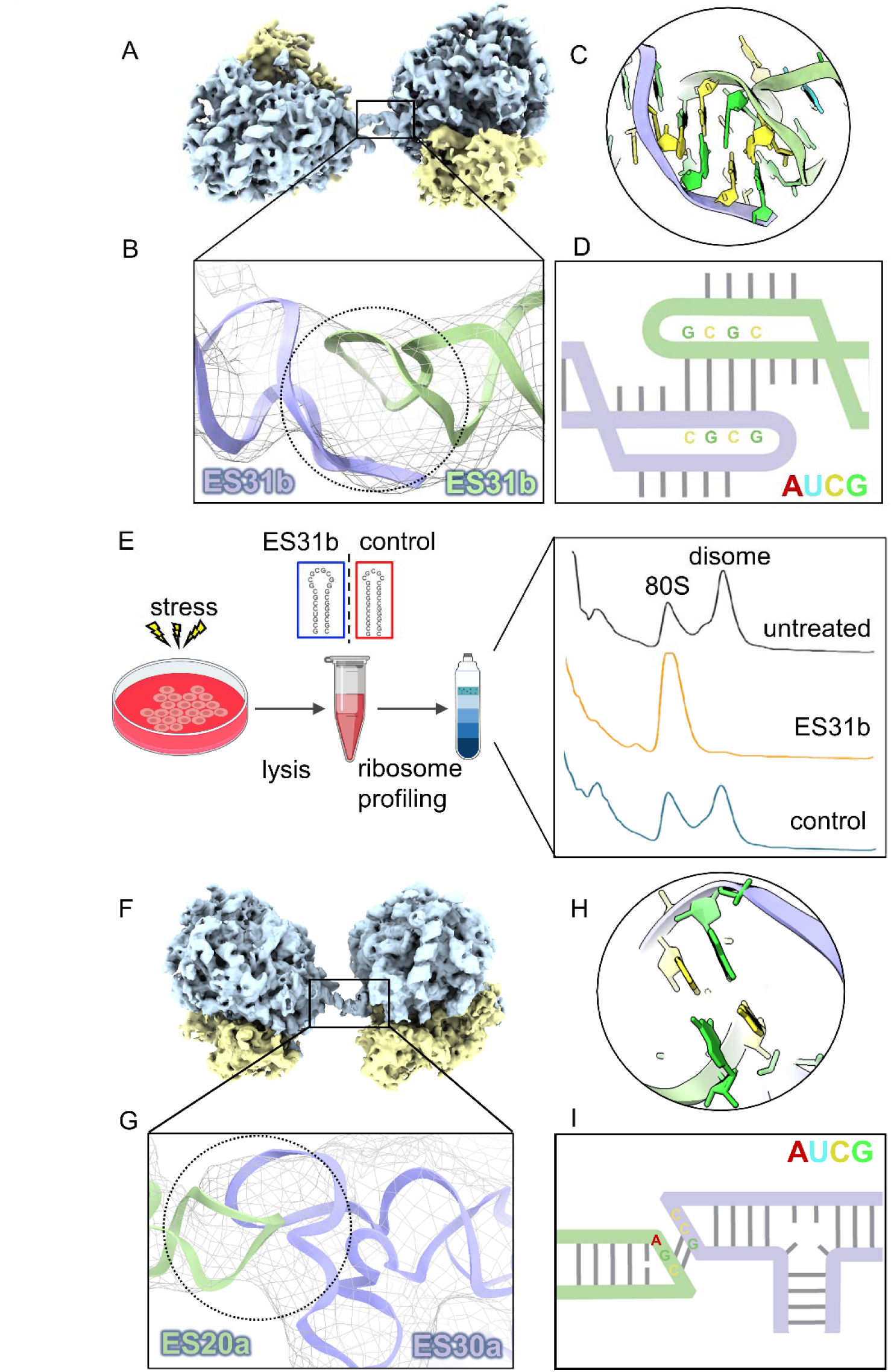
Idle ribosomes form dimers via rRNA expansion segment-mediated interactions. (A-D) Structural characterization of Disome 2. Overall architecture of Disome 2 is shown in A) with the large subunit in blue and the small subunit in yellow. A diagram of the atomic model fitted into the linkage region density is show in B). The inferred base-pairing interaction in the linkage region is shown in C). The secondary structure modeling of rRNA linkage region is shown in D). (E) Ribosome disomes are disrupted by addition of short hairpin RNA exhibiting the same sequence as ES31b. (F-I) Structural characterization of Disome 3, corresponding to (A-D).

Disome 2 is mediated by ES31 from both 80S ribosomes, resulting in a ribosome dimer exhibiting pseudo-C2 symmetry (Figure 4A). Structural characterization reveals ES31b adopts a hairpin conformation featuring a GC-rich loop. Sequence analysis indicates that this terminal loop potentially facilitates inter-ribosomal interaction via complementary base-pairing (Figure 4D).

To confirm this dimeric interface, we *in vitro* synthesized the isolated GC-rich RNA representing ribosomal ES31b sequence of rat neuronal cells that is expected to fold as a short (24 nucleotide) hairpin. A second RNA sequence that is predicted to form a similar hairpin structure, but with fewer GC residues in the GC-rich loop, was used as a control (Figure 4E). We then tested whether these two isolated RNA hairpin structures can compete with dimer-dimer interactions to disrupt the formation of disomes that formed in response to cell stress. The polysome profiles demonstrate that the disome peak dissociates in the presence of excess GC-rich RNA hairpin with ES31b sequence, whereas the shorter, control hairpin had no effect on ribosome dimer formation (Figure 4E). In agreement with this notion, structural analysis of the pooled disome peak revealed a strong enrichment of the Disome 2 conformation. Altogether, these results support that the dimeric interface is mediated by an RNA kissing loop regulated that can be outcompeted with an RNA hairpin comprised of the same ribosomal ES31b sequence. This observation supports the notion that the ES31b hairpin forms intermolecular Watson-Crick base pair interactions to form dimeric loops.

In contrast to the other two disome types and previously reported hibernation dimers with pseudo-C2 symmetry, Disome 3 is mediated by ES20a and ES30a of neighboring ribosomes. Steric hindrance prevents the formation of higher-order oligomers (Figure 4F-I). ES30a is part of the L1 stalk, whose flexibility is critical for translation elongation ^36^. Therefore, the formation of such a dimer would be incompatible with translation activity. Notably, as mentioned above, eIF5A in the idle state 80S ribosome could stabilize the L1 stalk by interaction with uL1, potentially enhancing ES30a-ES20a interaction.

Ribophagy, the selective autophagy of ribosomes, is critically regulated by ubiquitination dynamics of ribosomal proteins, particularly at uL23/RPL25 in yeast under starvation conditions ^37^. Strikingly, our structural mapping reveals that uL23 localizes proximally to both ES30a and ES31b (Figure S7C-D), suggesting that the formation of Disome 2 and Disome 3 may sterically hinder recognition of uL23 by ubiquitylation-related enzymes. This spatial arrangement raises the possibility that translationally inactive ribosomes may be less accessible to autophagic recognition, although direct experimental validation of this hypothesis remains pending.

The sequences of rRNA expansion segments exhibit high diversity, while their functions remain largely unknown. To investigate whether such a translational regulatory mechanism is potentially evolutionarily conserved across taxa, we performed sequence analysis of rRNA expansion segments located at interface regions across diverse species. This analysis revealed distinct patterns: Disome 3 formation appears to be restricted to vertebrate lineages examined in this analysis. Conversely, Disome 2 exhibits broader conservation across different species, and shows highly consistent sequence features particularly in humans, suggesting that it may represent a more evolutionarily conserved form of idle ribosomes (Figure S8).

### Reversible ribosome dimerization in response to multiple stress conditions

In bacteria, the formation of 100S ribosome dimers serves as a protective mechanism under various stress conditions. We hypothesized that rRNA expansion segment-mediated ribosome dimerization may represent a universal stress response mechanism beyond the puromycin treatment discussed above. To investigate this, we treated C6 glioma cells with puromycin, Krebs-Ringer Bicarbonate (KRB) buffer to induce amino acid deficiency or with administration of cyclopiazonic acid (CPA), a reversible endoplasmic reticulum inducer. Ribosome profiling revealed that, like puromycin treatment, these conditions resulted in accumulation of disomes, which coincided with a decrease in translating polysomes (Figure S9A). To determine whether this process is reversible, we followed administration of CPA or puromycin with a washout to remove these reagents from the media. Ribosome profiling revealed that washout of CPA or puromycin reduced the population of disomes (Figure S9B-C), while restoring levels of translating ribosomes following 2-6 hours of washout (Figure S9B). These data indicate that disome formation in C6 glioma cells represents a reversible hibernating ribosome response to different inducers of cellular stress. which may modulate the translation activity in response to diverse cellular stresses.

## Discussion

Localized mRNA translation plays a critical role in neuronal development and function ^38^. A widely accepted hypothesis suggests that mRNA is transported from the soma to distal neuronal sites in either repressed or stalled polysome states. In agreement with this concept, populations of purified neuronal ribosomes include a significant proportion stalled in a hybrid-state, as indicated by their ability to retain puromycin-modified peptides ^18^. However, in our work, while the majority of ribosomes are engaged in translation elongation, topological analysis has identified only a subset of stalled-collided disomes, which constituted only 1.2% of the total ribosome population. These disomes are characterized by GCN1 recruitment and hybrid-state of the stalled ribosomes (Figure S10). Notably, the population of these structures is insufficient to form clusters in the soma region, irrespective of puromycin treatment. Given these observations *in situ*, we propose that certain translation elongation ribosome complexes may exhibit high fragility, such that even mild purification procedures disrupt their structural integrity. This underscores the necessity of *in situ* structural analysis to accurately capture their native state.

Puromycin and its derivatives are extensively employed in cell biology due to their ability to non-selectively incorporate into nascent peptides ^39^. On the one hand, our findings reveal that membrane-bound ribosomes display heightened susceptibility to puromycin treatment compared to cytosolic ribosomes (Figure S11). On the other hand, in addition to the rapid accumulation of idle ribosomes and the formation of potentially hibernating disomes, our research provides the first evidence of eIF5A’s role in modulating puromycylation efficiency. These observations suggest that puromycin may exhibit preferential targeting and that its activity is regulated by other cellular factors, leading to complex cellular outcomes. Consequently, caution is warranted when interpreting results from puromycin-based assays in cell biology studies.

Expansion segments are evolutionary insertions of sequence blocks that contribute to the larger ribosome size in eukaryotes. However, only a limited number of expansion segments have been functionally characterized ^40–43^. Our data suggest that expansion segments may play a role in the formation of hibernating ribosome dimers, a process mediated by protein factors in prokaryotes. Intriguingly, recent work from our group has identified a non-translating disome whose formation is mediated by ES27 and the protein factor EBPl ^44^, further emphasizing the broader functional significance of expansion segments in the formation of non-translating disomes. Beyond direct translation inhibition by puromycin, we demonstrate that diverse stress conditions, including nutrient deprivation, unfolded protein response, and ER stress, can also induce disome assembly. The reversible nature of this dimerization, coupled with its negative correlation with translation activity, supports a role for ribosome dimerization as a general regulatory mechanism to modulate protein synthesis.

In conclusion, through the application of *in situ* cryo-electron tomography (cryo-ET) and topological analysis, we demonstrate that puromycin treatment significantly alters the translation landscape at both the individual 80S ribosome and polyribosome levels. Run-off idle ribosomes are further clustered to form eukaryotic hibernating ribosome dimers, a process mediated by expansion segments. This highlights a novel role for rRNA expansion segments in the cellular stress response. However, our study does not establish a definitive causal relationship between dimer formation and translation regulation, partly due to the technical challenges associated with rDNA gene editing. Additionally, given the inherent flexibility of RNA structures and the complexity of the crowded cellular environment, future research should focus on inter-molecular interactions rather than individual molecules to fully elucidate the mechanisms underlying translation regulation.

## Methods

### Cell Culture

Rat hippocampal neurons were isolated and cultured as described previously ^45^. After cutting the rat brain tissue into small pieces, the samples were digested with trypsin for 15 minutes. Following digestion, an equal volume of DMEM-F12 medium (Thermo Fisher #11320033) containing 10% fetal bovine serum was added to dissociate the tissue and release each single cell. The mixture was left undisturbed for 2 minutes to allow cell sedimentation, after which the supernatant was transferred to a new tube and centrifuged at 500 g for 2 minutes. The cells were then resuspended with fresh DMEM-F12 medium containing 10% FBS and seeded into culture dishes pre-coated with poly-D-lysine (Sigma-Aldrich #P7280). After 2 hours, a neuronal culture medium consisting of B27 (Thermo Fisher #17504044), GlutaMAX (Thermo Fisher #35050061), and Neurobasal medium (Thermo Fisher #21103049) was added. 24 hours later, 2 μM cytosine arabinoside (Ara-C) (Sigma-Aldrich #C3350000) was added to the medium to remove glial cells. The cells were grown at 37°C and 5% CO_2_. The medium was refreshed every 48 hours, with half of the medium being replaced.

C6 rat glioma cells (ATCC, CCL-107) were cultured in high-glucose Dulbecco’s Modified Eagle’s Medium (DMEM, Gibco #11960044) supplemented with 10% fetal bovine serum (FBS), 2 mM L-glutamine, and antibiotics (penicillin and streptomycin, Gibco #15070063). Cells were maintained at 37°C in a humidified atmosphere with 5% CO_2_. For amino acid starvation experiments, cells were incubated in Krebs-Ringer Bicarbonate Buffer (KRB, Merck #K4002) supplemented with 10% dialyzed FBS. Control cells were simultaneously transferred to fresh DMEM supplemented with 10% dialyzed FBS. For drug treatments, cells were exposed to cyclopiazonic acid (CPA, Tocris #1235) at a final concentration of 200 μM or puromycin (Gibco #A1113803) at 275 μM. For inhibitor washout experiments, after the designated treatment duration, the medium containing inhibitors was removed, cells were washed twice with warm Dulbecco’s Phosphate-Buffered Saline (dPBS, Gibco #14040133), and fresh warm DMEM was added.

### Ribosome Profiling

C6 rat glioma cells were prepared as described above. Translation was halted by treating cells with cycloheximide (CHX, Merck #C7698) at a final concentration of 100 μg/ml for 3 minutes at 37°C. Cells were then washed twice with PBS containing 100 μg/ml CHX, scraped, and pelleted by centrifugation at 3,500 rpm for 10 minutes at 4°C. Cell pellets were resuspended in 500 μl of lysis buffer (10 mM HEPES-KOH pH 7.4, 2.5 mM MgCl2, 100 mM KCl, 0.25% Nonidet NP-40, 200 units/ml RNaseOUT [Invitrogen]) and incubated on ice for 20 minutes. Lysates were homogenized by passing 15 times through a 23-gauge needle and cleared of debris by centrifugation at 12,000 rpm for 10 minutes at 4°C. The supernatant was collected, and RNA concentration was determined by absorbance measurements at 260 nm using a BioTek Take3 reader with a Microvolume Plate (Agilent).

Polysomes were separated using 10–50% sucrose gradients (10 mM HEPES-KOH pH 7.4, 2.5 mM MgCl_2_, 100 mM KCl) prepared in an SW28 rotor (Beckman) centrifuged at 17,000 rpm for 15 hours or an SW41Ti rotor centrifuged at 40,000 rpm for 2 hours at 4°C. Finally, the resulting sucrose gradient solution was analyzed using a Biocomp instrument.

### Sample preparation

For cryo-EM analysis, R2/1 holey carbon gold grids (T10012 Lacey Formvar Carbon Coated G-Grids, Beijing XXBR Technology) were coated with an additional carbon layer (∼20 nm thick) using a carbon evaporator (ACE 600) and neuronal cells were seeded at a density of 3 x 10^6^ cells/ml following 6 days of culture. For stress condition, cells were incubated with 275 μM puromycin for 10 minutes before vitrification. Afterwards, cell culture medium was replaced with medium containing 10% glycerol. Then grids were blotted for 10 seconds using filter paper and immediately plunged into liquid ethane based on a Vitroboot (FEI). Grids were stored in liquid nitrogen until further processing.

### Lamella preparation

The lamellas were prepared in the regions of neuronal soma using a cryo-FIB-SEM system (Thermo Fisher Scientific #Aquilos^TM^). In brief, grids were subjected to platinum sputtering (15 seconds) followed by a gas injection system (GIS) treatment to deposit an additional organomtallic platinum protective layer for 30 seconds. Samples were tilted to angle of 12° for following milling with an ion beam of 30 kV. In the initial milling process, we used 0.5 nA for preliminary thinning, and then gradually adopted smaller beam current for the fine process. The final thickness of the lamellas was about 150 nm. SEM imaging was used to monitor the milling progress.

### Cryo-ET data collection

The samples were examined at liquid nitrogen temperature on a Titan Krios (Thermo Fisher Scientific) equipped with a K3 summit direct electron detector and Bioquantum energy filter (Gatan). The microscope was operated at an accelerated voltage of 300 kV and 20 eV slit. Images were recorded in movies of 10 frames at a target defocus of 2 to 5 μm and pixel size of 1.37 Å. Tilt series were required from −40° to +60° with an angular increment of 2° using a grouped dose-symmetric tilt scheme in SerialEM (3.7) ^46^. The cumulative dose of a series did not exceed 110 e^−^/Å.

### Tomogram reconstruction and particle picking

Each tilt image frames were motion-corrected using MotionCorr2 ^47^. After that, tilt series were aligned using fiducial-free patch tracking strategy, and the tomograms were reconstructed by weighted back-projection using the IMOD (4.11.3) ^48^ at a pixel size of 8.22 Å (6x binned). CTF estimation for individual projection images as well as entire tile series was performed in Warp (1.1.0) ^49^. Ribsome particle coordinates were determined using deepFinder (0.0.1) ^50^. And false positive particles were manually filtered.

### Subtomogram analysis

The subtomogram analysis workflow is represented in Figure S2. Specifically, the determined positions of particles were used to extract subtomograms and corresponding CTF volumes at a pixel size of 5.48 Å (4x binned) using Warp ^49^. Those extracted subtomograms were 3D classified with volume alignment against a low pass filter 80S ribosome map (EMDB-33077) as template in RELION (3.1.4) ^51^ to exclude false positive. The remaining subtomograms were further refined in RELION and subjected to an iterative refinement in M (1.1.0) ^49^, resulting in a ribosome density map with an overall 4.8 Å resolution. The particles were re-extracted in Warp at pixel size of 2.74 Å (2x binned). Bin2 subtomograms were refined in RELION with a mask covering the entire 80S ribosome. After that, 3D classification was performed with local volume alignment with a mask covering the LSU to separated LSU particles. A second round of classification was performed with a mask covering the SSU, resulting in rotated and non-rotated 80S ribosomes groups. Then masks covering the tRNA or elongation factor binding sites were adopted to classified ribosomes at different translation states. The resulting conformations were further subjected into 3D classification with different parameters to ensure no new conformation detected. All classes were finally subjected to refinement in RELION to improve the resolution.

### Cryo-EM data acquisition and processing

Data acquisition was performed using a Titan Krios cryo-EM (Thermo Fisher Scientific) operating at 300 kV, equipped with an energy filter (Gatan) and a Falcon 4i direct electron detector. Imaging was conducted with a pixel size of 1.21 Å. For single-particle data collection, a multi-shot pattern was employed at a single tilt angle of -12 or 12 degrees to compensate for the lamellae sectioning angle. Defocus values ranged from -0.8 to -1.6 μm, and each shot received a total dose of 50 e/Å². Data acquisition was controlled using SerialEM software. A total of 6,904 movies were collected for the puromycin-treated samples. Motion correction was applied to each image frame using MotionCorr2, and the CTF values were determined with CTFFind3 ^52^. Afterwards, particles were located using GisSPA, and the initial reference was derived from the ribosome map (EMD:23500) with small subunit manually cropped. Several rounds of 3D classification were applied to filter false-positive particles in RELION, followed by 3D refinement, CTF refinement and Bayesian polishing to further improve the resolution. The large subunit of resulting high-resolution 80S ribosome map was adopted to re-localized the ribosome particles by GisSPA, following the similar processing steps mentioned above, resulting 117,690 80S particles. For idle ribosome classification, an initial 3D classification was performed using a mask covering the tRNA binding sites to separate ribosomes engaged in translation elongation. Ribosomes exhibiting eEF2-like density occupancy in the A-site and empty P-site, representing 72.1% of the total population, were kept for subsequent analysis. A secondary 3D classification was then conducted using a tighter mask localized to the eIF5A binding region to further separate idle states subpopulations. This revealed two distinct particle subsets: 41.5% (48,855 particles) of particles contained eEF2, SERBP1 and eIF5A, while 15.4% (35,924 particles) with only eEF2 and SERBP1. These two idle states particles underwent iterative processing to optimize resolution, including 3D refinement, CTF refinement, and Bayesian polishing in RELION. Initial structures of factors (eIF5A, SERBP1, eEF2) were modeled based on the published model (PDB ID:6Z6M), following rigid-body fitting into cryo-EM maps using ChimeraX (1.3) ^53^. Models were manually adjusted in Coot (0.9.7) ^54^ before being imported into ISOLDE (1.6) ^55^ within ChimeraX to improve dihedral angles and rotamer fitting.

### Neighboring ribosome topology characterization and visualization

All 80S ribosomes were subjected to NEMO-TOC for neighboring ribosome topology analysis with default parameters. The identified clusters were further categorized into five distinct groups based on structural features: (1) cytosolic helical polysomes (cyto-helical), (2) membrane-bound polysomes (membrane-bound), and three dimer subtypes (Disome 1/2/3) differentiated by inter-ribosomal rRNA contact interface. Besides, stalled-collided dimers were identified by optimized NEMO-TOC detection threshold, and structurally validated by GCN1-like density attached in cryo-EM reconstructions. For structural visualization, representative neighboring ribosome topologies were reconstructed through placing adjacent ribosome maps using determined rotation and translocation vectors determined by NEMO-TOC, with corresponding composite maps presented in Figure3A.

### rRNA secondary structure analysis

Secondary structure annotations of LSU across multiple species were retrieved from the RNA central database ^56^ for comparative analysis.

### RNA interference

An appropriate number of C6 rat glioma cells (1.0×10^5 cells each well) were seeded into a 12-well plate and culture overnight. When cell density reaches approximately 50% confluency, siRNA transfection was performed. The transfection processing was conducted according to the method described by Karina et al. ^57^ using Opti-MEM medium and Lipofectamine® RNAiMAX transfection reagent. The siRNA sequences (5’-3’) designed was: (sense) GAAGAUAUCUGCCCGUCAA and (antisense) UUGACGGGCAGAUAUCUUCTT for knockdown; (sense) UUCUCCGAACGUGUCACGUTT and (antisense) ACGUGACACGUUCGGAGAATT for negative control. After 12 hours, the medium was replaced with fresh culture medium, followed by medium replacement every 24 hours. Approximately 60 hours post-transfection, cells were passaged into 12-well plates (1.8×10^5 cells each well). After 12 hours, the second round of siRNA transfection were performed. The experimental procedures were performed 24 hours later.

### SDS PAGE and Western Blotting

C6 rat glioma cells were washed twice and scraped in ice-cold PBS supplemented with M5 Phosphatase Inhibitor Cocktail (Roche). The solution was then centrifuged at 2000 rpm for 5 minutes at 4°C to remove the supernatant. After adding cell lysis buffer RIPA (Beyotime #P0013K) supplemented with M5 Phosphatase Inhibitor Cocktail at 4°C for 10 minutes, the lysate was centrifuged at 12,000 rpm for 10 minutes to collect the supernatant. The supernatant was diluted in 5X SDS-PAGE sample loading buffer (Beyotime #P0015) and heated at 100°C for 10 minutes. After that, the sample was loaded onto 10% polyacrylamide gels. Afterwards, samples were transferred onto 0.45 µm PVDF membranes (MERCK #IPVH00010) by wet transfer and blocked with 5% no-fat milk in TBS-T (20 mM Tris-HCl with pH 7.5, 137 mM NaCl, 0.1% Tween 20). The membrane was incubated overnight at 4℃ with primary antibodies (anti-puromycin AB 2619605) (1:1,000), followed by HRP-conjugated secondary antibodies (ThermoFisher #31430, #31460) incubation at room temperature for 1 hour and detected by chemiluminescence (Tanon 5200). After each antibody incubation, the membrane was washed three times with TBST for 15 minutes. In some cases, the blots were stripped (ZmTech Scientific #S208070) and reblotted with an anti-eIF5a antibody (Cell Signaling #2217) or with anti-GADPH antibody (Proteintech #14583-1-AP).

### Visualization

Membranes were segmented using Membrain software ^58^. All figures were prepared by ChimeraX ^53^ and Python (3.7).

## Acknowledgments

We are grateful to the Cryo-EM Platform of Peking University and Changping Laboratory, the High-Performance Computing Platform of Peking University for the support on data collection and computation, the National Centre for Protein Sciences at Peking University for technical assistance. This work is funded by the National Key Research and Development Program of China (#2024YFA1802800 to Q.G.), the National Natural Science Foundation of China (#32371191 to Q.G.), the Beijing Natural Science Foundation (JQ24031 to Q.G.). Q.G. is supported by Changping Laboratory, the SLS-Qidong Innovation Fund and the Li Ge-Zhao Ning Life Science Youth Research Foundation.

---

**Figure S1.**
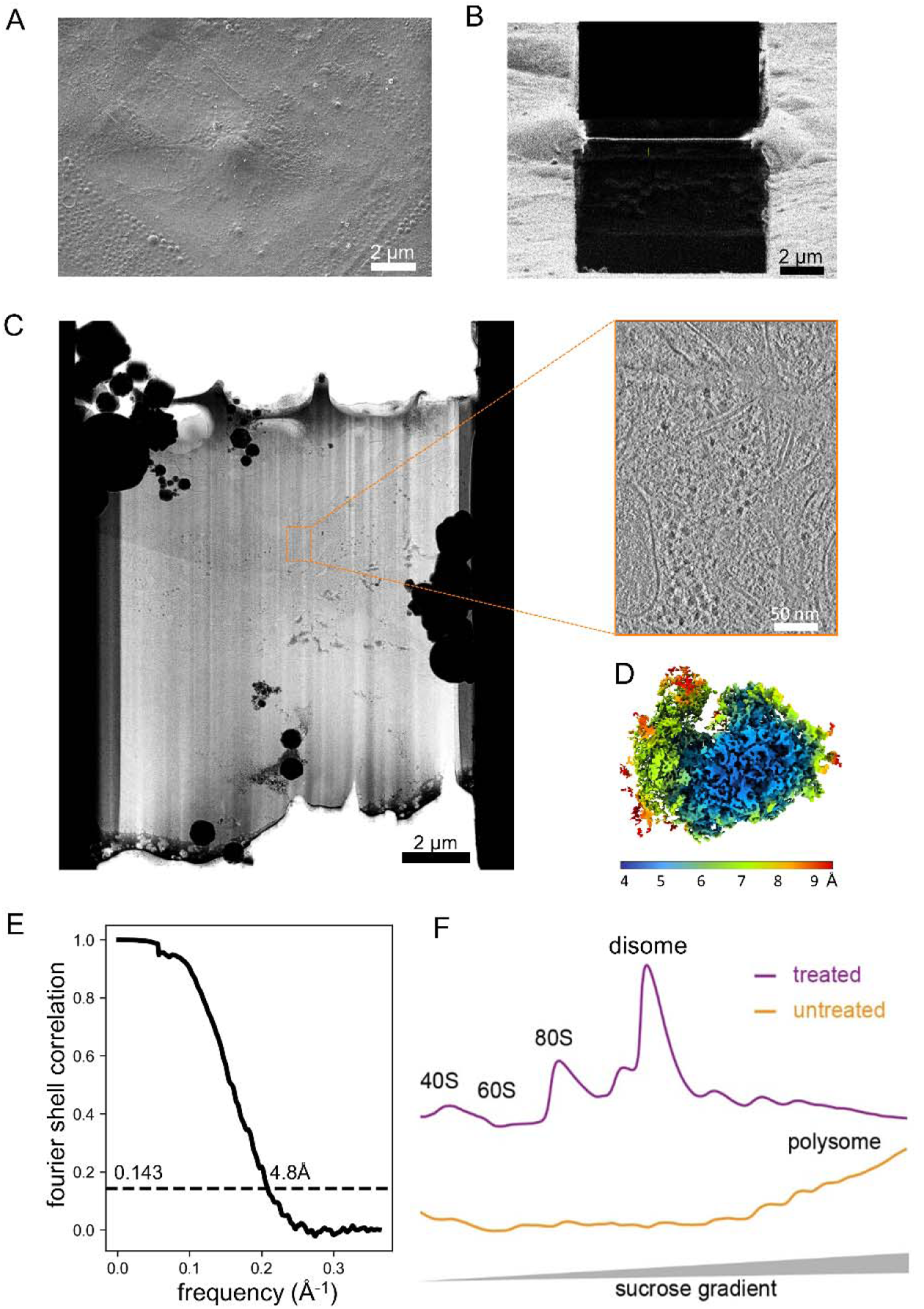
Experimental pipeline from sample preparation to structural analysis. (A) SEM image showing neuronal cells cultured on EM grids. (B-C) The cellular lamella was prepared using cryo-FIB with a thickness of < 200 nm. (D) The subtomogram averaging map of the ribosome color-coded by local resolution. (E) The FSC curve assessing reconstructed map’s resolution. (F) Ribosome profiling results using sucrose gradients: control (orange) vs. puromycin-treated (purple).

**Figure S2.**
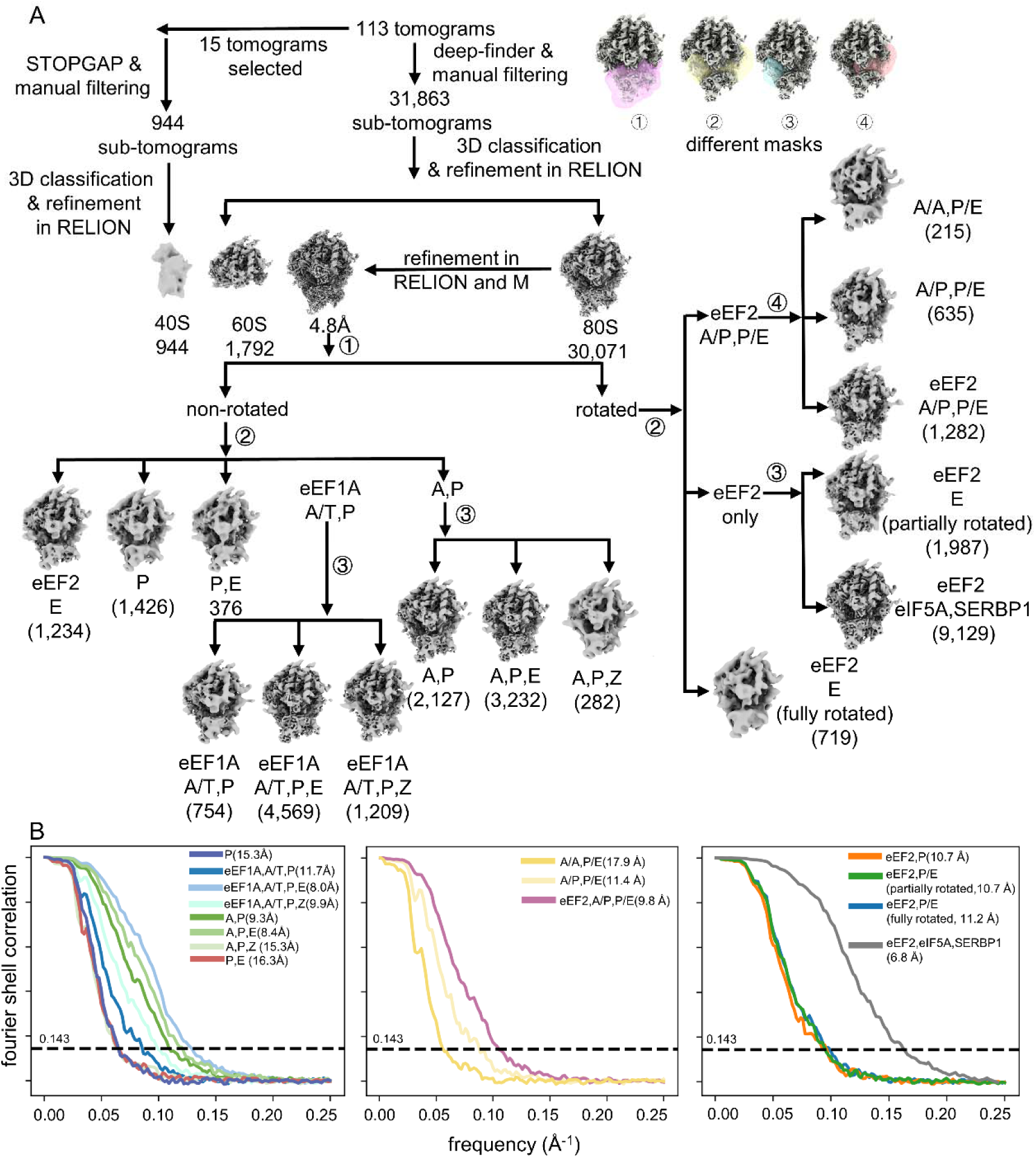
Ribosome classification pipeline in cryo-ET datasets. (A) Region-specific masks were applied for ribosome conformation classification. (B) FSC analysis of different classified ribosome conformations.

**Figure S3.**
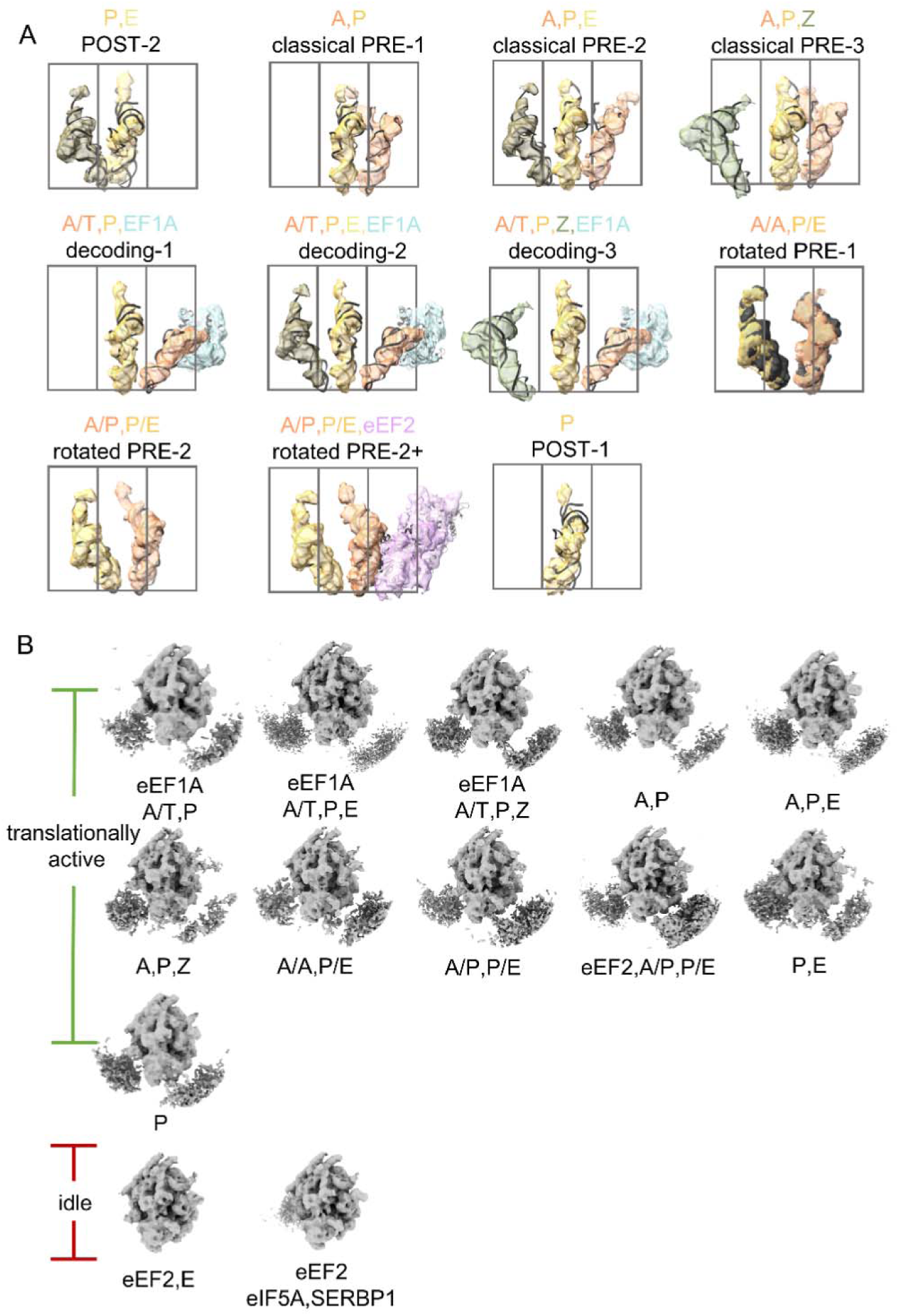
The assigned translation states of ribosomes. (A) The tRNA occupancy and elongation factors binding patterns across each translationally active ribosome conformation. (B) The neighborhood density of individual ribosome translation state.

**Figure S4.**
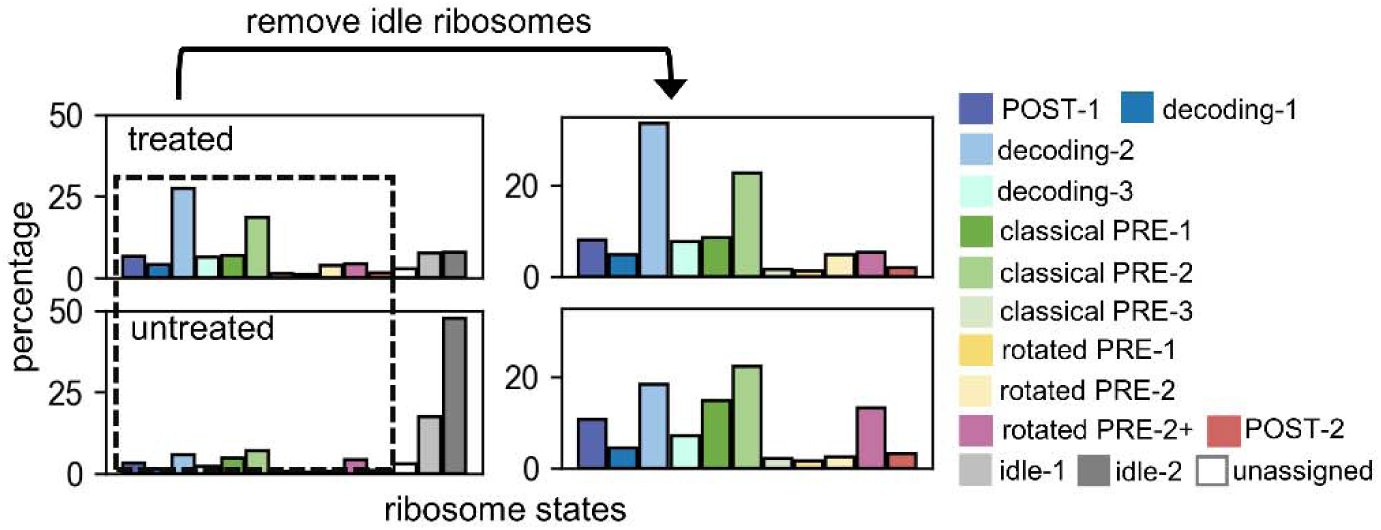
**Comparative analysis of ribosome translation states populations: untreated vs. puromycin-treated with/without idle ribosomes inclusion.**

**Figure S5.**
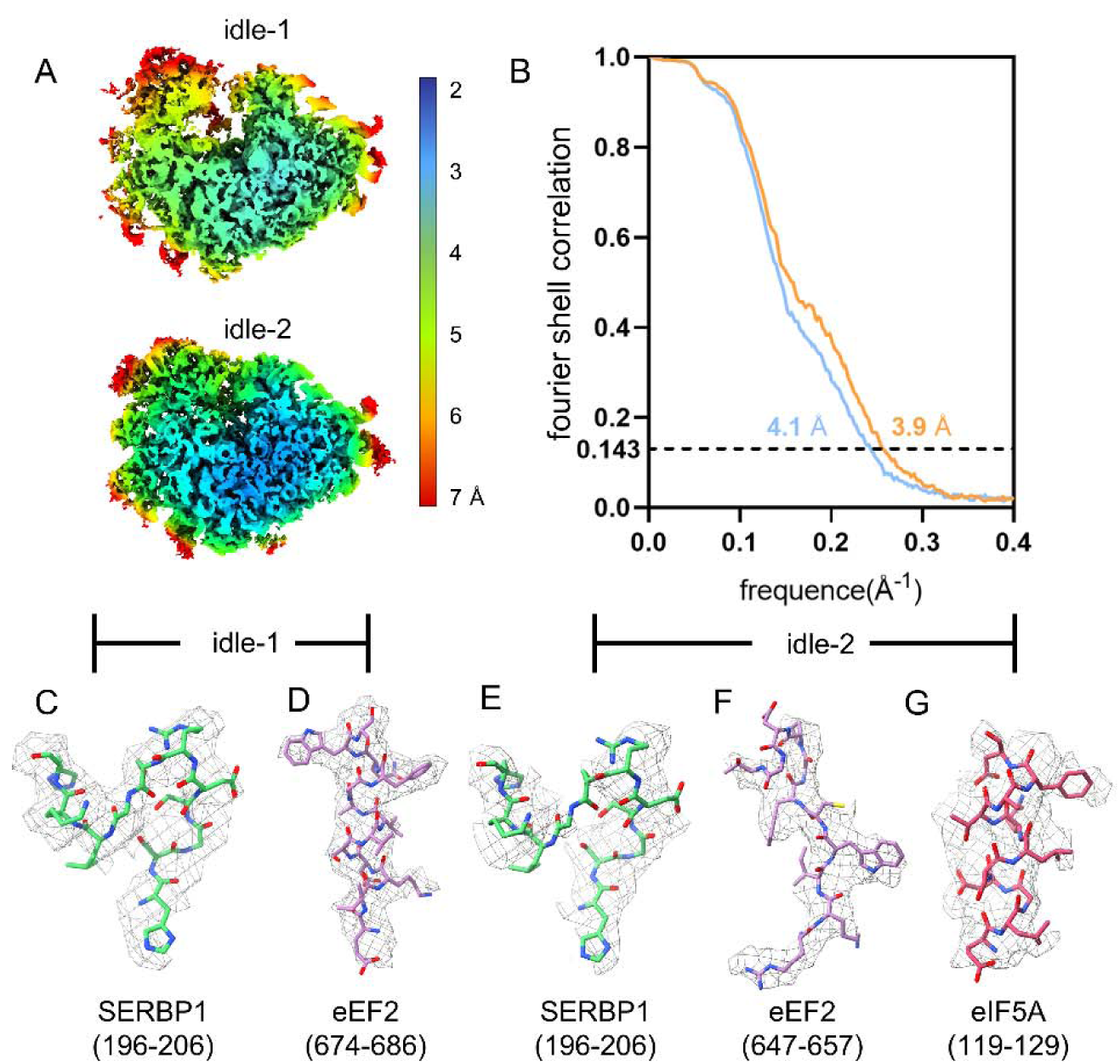
Cryo-EM analysis of idle ribosomes using GisSPA. (A-B) Local resolution maps and FSC assessment for ribosomes of two different idle states. (C-D) Atomic model validation of idle-1 ribosome with density fitting: SERBP1 and eEF2. (E-G) Atomic model validation of idle-2 ribosome with density fitting: SERBP1, eEF2 and eIF5A.

**Figure S6.**
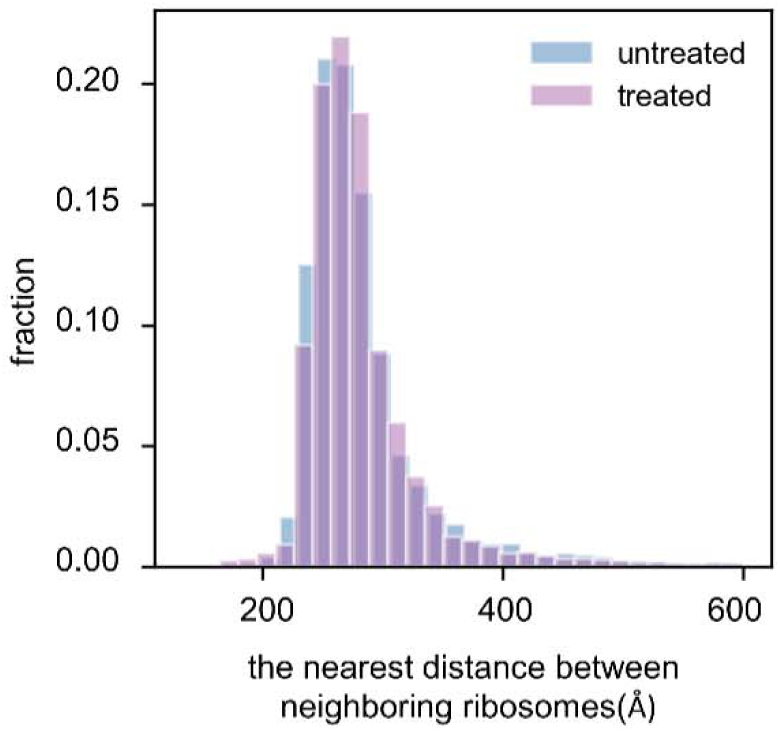
The distribution of neighboring ribosomes distances. Puromycin-treated ones are in purple and untreated ones are in blue.

**Figure S7.**
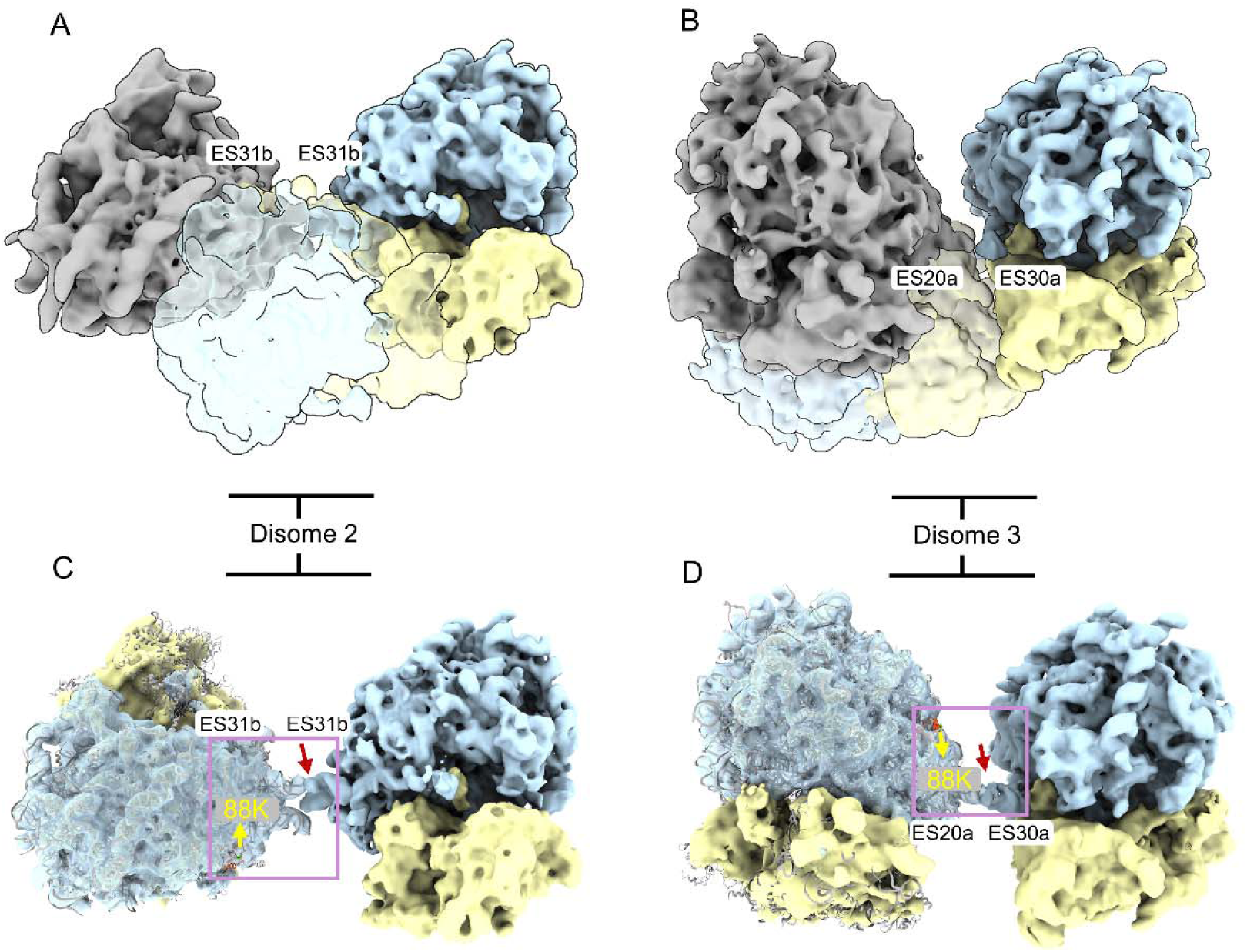
Structural characteristics of the two hibernating disomes. (A-B) A comparative structural analysis of Disome 2 (A) and Disome 3 (B), depicted in gray, is presented. These disomes are compared against the previously characterized collided-stalled disome (EMDB-4427) by aligning to the stalled ribosome. Large ribosomal subunits are colored in blue, while small subunits are highlighted in yellow for clarity. (C-D) The reported ribophagy-associated ubiquitination site (lysine 88, indicated by a yellow arrow) ^37^ in uL23 (red) and the interface region of the dimers in this work (marked with a red arrow) are proximal to each other.

**Figure S8.**
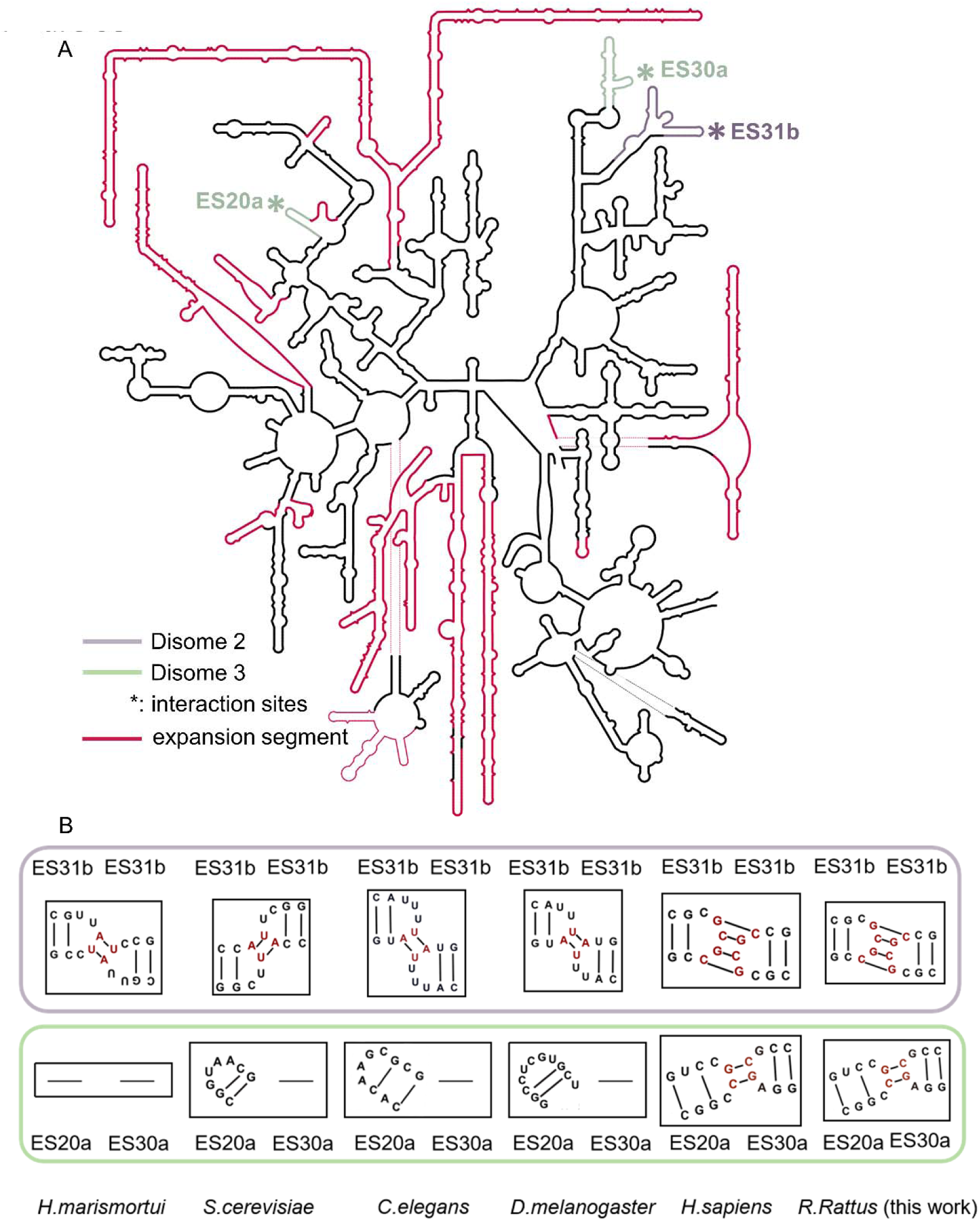
The rRNA sequence analysis at disome interfaces. (A) The secondary structure of rat 28S rRNA is illustrated, with expansion segments highlighted in red. The interface regions associated with Disome 2 (labeled in purple) and Disome 3 (labeled in green) are denoted by asterisks. (B) The inferred base-pairings within these interface regions of Disome 2 (purple boxed) and Disome 3 (green boxed) across different species. Noting in some species, specific expansion segments are absent and are denoted with ‘-‘.

**Figure S9.**
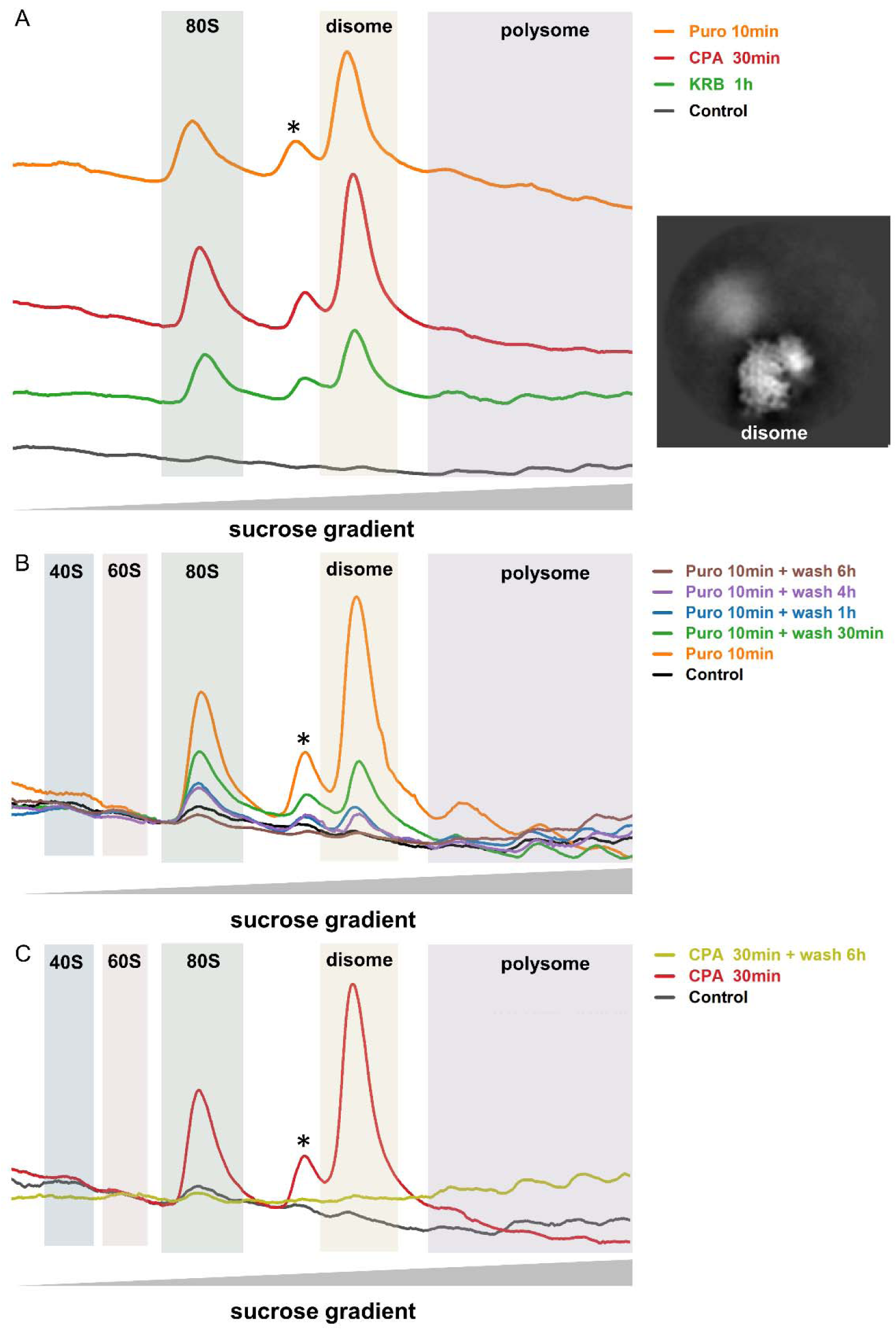
Disome formation in response to different stress stimuli in C6 glioma cells is reversible. (A) Ribosome profiling shows that ribosome disomes increase in response to puromycin administration, CPA administration, or amino acid starvation (KRB media). The right panel shows the 2D projection of the purified disomes, as determined by the single-particle cryo-EM. (B) Ribosome disomes formed under puromycin administration is reversible following washout. (C) Ribosome disomes formed under CPA administration is reversible following washout. The * peak is indicative of a putative 60S-80S disome.

**Figure S10.**
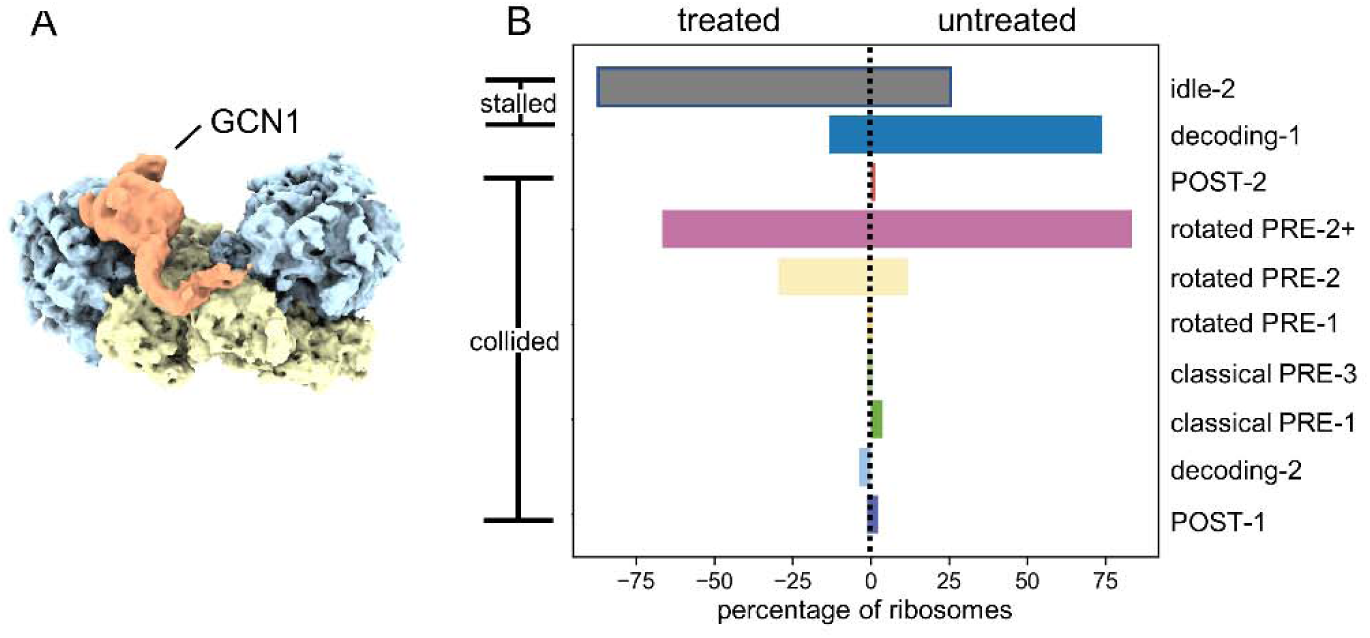
In situ analysis of ribosome collision. (A) In situ subtomogram average of GCN1-bound (orange) collided-stalled disomes. (B) The ribosome translation states distribution of stalled or collided ribosomes in the untreated and puromycin-treated group.

**Figure S11.**
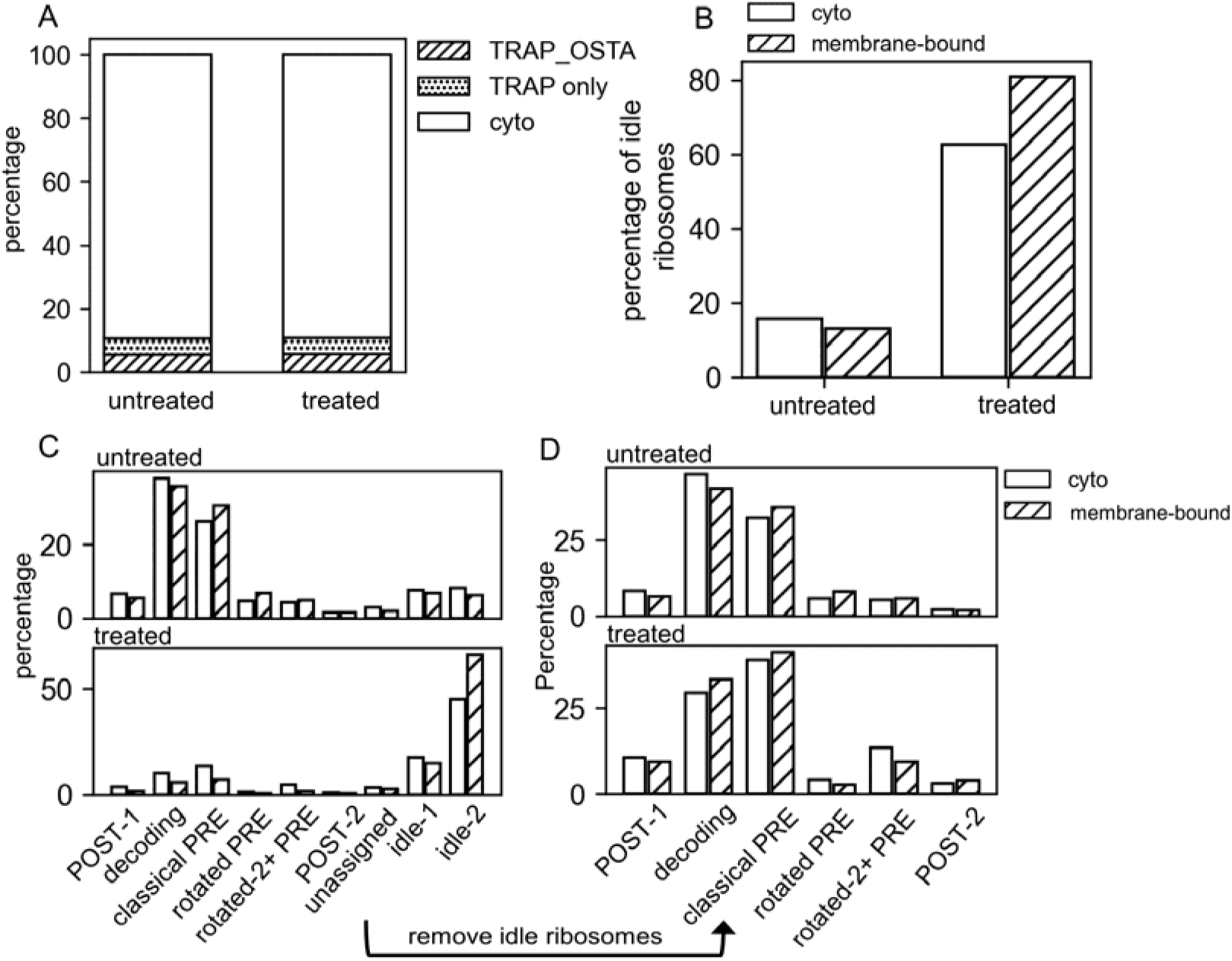
Comparison between cytoplasmic and membrane-bound ribosomes. (A) The population of cytoplasmic ribosomes, membrane-bound ribosome with/without OSTA in untreated or puromycin-treated group. (B) The population of idle ribosomes in the cytoplasm or bound to the membrane in untreated and puromycin-treated group. (C-D) Translation states distribution of ribosomes in untreated and puromycin-treated group with (C) and without (D) the inclusion of idle ribosomes.

**Table S1.**
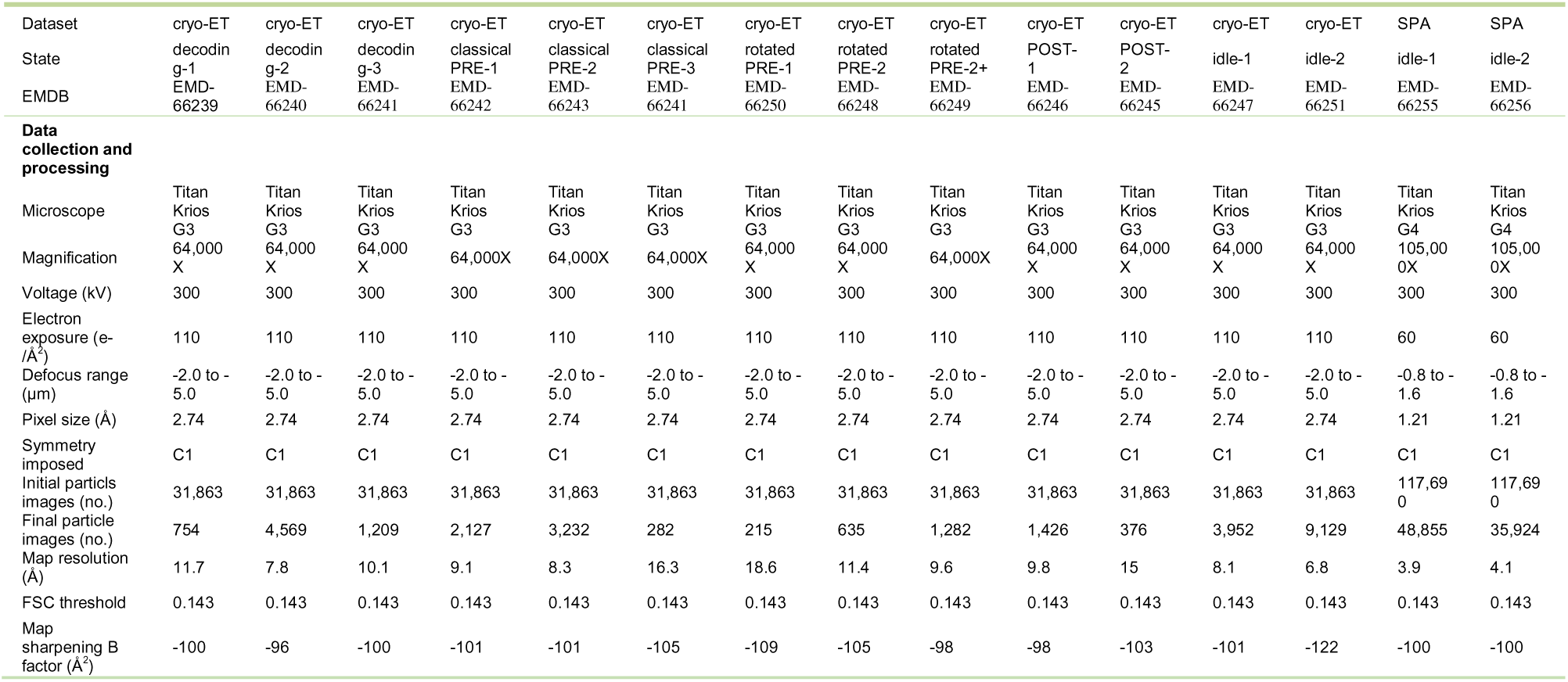
Single particle cryo-EM and cryo-ET data collection.

| Dataset | cryo-ET | cryo-ET | cryo-ET | cryo-ET | cryo-ET | cryo-ET | cryo-ET | cryo-ET | cryo-ET | cryo-ET | cryo-ET | cryo-ET | cryo-ET | SPA | SPA |
| --- | --- | --- | --- | --- | --- | --- | --- | --- | --- | --- | --- | --- | --- | --- | --- |
| State | decodin<br>g-1 | decodin<br>g-2 | decodin<br>g-3 | classical<br>PRE-1 | classical<br>PRE-2 | classical<br>PRE-3 | rotated<br>PRE-1 | rotated<br>PRE-2 | rotated<br>PRE-2+ | POST-<br>1 | POST-<br>2 | idle-1 | idle-2 | idle-1 | idle-2 |
| EMDB | EMD-66239 | EMD-66240 | EMD-66241 | EMD-66242 | EMD-66243 | EMD-66241 | EMD-66250 | EMD-66248 | EMD-66249 | EMD-66246 | EMD-66245 | EMD-66247 | EMD-66251 | EMD-66255 | EMD-66256 |
| **Data collection and processing** |  |  |  |  |  |  |  |  |  |  |  |  |  |  |  |
| Microscope | Titan<br>Krios<br>G3 | Titan<br>Krios<br>G3 | Titan<br>Krios<br>G3 | Titan<br>Krios<br>G3 | Titan<br>Krios<br>G3 | Titan<br>Krios<br>G3 | Titan<br>Krios<br>G3 | Titan<br>Krios<br>G3 | Titan<br>Krios<br>G3 | Titan<br>Krios<br>G3 | Titan<br>Krios<br>G3 | Titan<br>Krios<br>G3 | Titan<br>Krios<br>G3 | Titan<br>Krios<br>G4 | Titan<br>Krios<br>G4 |
| Magnification | 64,000<br>X | 64,000<br>X | 64,000<br>X | 64,000X | 64,000X | 64,000X | 64,000<br>X | 64,000<br>X | 64,000X | 64,000<br>X | 64,000<br>X | 64,000<br>X | 64,000<br>X | 105,00<br>0X | 105,00<br>0X |
| Voltage (kV) | 300 | 300 | 300 | 300 | 300 | 300 | 300 | 300 | 300 | 300 | 300 | 300 | 300 | 300 | 300 |
| Electron exposure (e-/Å<sup>2</sup>) | 110 | 110 | 110 | 110 | 110 | 110 | 110 | 110 | 110 | 110 | 110 | 110 | 110 | 60 | 60 |
| Defocus range (µm) | -2.0 to -<br>5.0 | -2.0 to -<br>5.0 | -2.0 to -<br>5.0 | -2.0 to -<br>5.0 | -2.0 to -<br>5.0 | -2.0 to -<br>5.0 | -2.0 to -<br>5.0 | -2.0 to -<br>5.0 | -2.0 to -<br>5.0 | -2.0 to -<br>5.0 | -2.0 to -<br>5.0 | -2.0 to -<br>5.0 | -2.0 to -<br>5.0 | -0.8 to -<br>1.6 | -0.8 to -<br>1.6 |
| Pixel size (Å) | 2.74 | 2.74 | 2.74 | 2.74 | 2.74 | 2.74 | 2.74 | 2.74 | 2.74 | 2.74 | 2.74 | 2.74 | 2.74 | 1.21 | 1.21 |
| Symmetry imposed | C1 | C1 | C1 | C1 | C1 | C1 | C1 | C1 | C1 | C1 | C1 | C1 | C1 | C1 | C1 |
| Initial particles images (no.) | 31,863 | 31,863 | 31,863 | 31,863 | 31,863 | 31,863 | 31,863 | 31,863 | 31,863 | 31,863 | 31,863 | 31,863 | 31,863 | 117,69<br>0 | 117,69<br>0 |
| Final particle images (no.) | 754 | 4,569 | 1,209 | 2,127 | 3,232 | 282 | 215 | 635 | 1,282 | 1,426 | 376 | 3,952 | 9,129 | 48,855 | 35,924 |
| Map resolution (Å) | 11.7 | 7.8 | 10.1 | 9.1 | 8.3 | 16.3 | 18.6 | 11.4 | 9.6 | 9.8 | 15 | 8.1 | 6.8 | 3.9 | 4.1 |
| FSC threshold | 0.143 | 0.143 | 0.143 | 0.143 | 0.143 | 0.143 | 0.143 | 0.143 | 0.143 | 0.143 | 0.143 | 0.143 | 0.143 | 0.143 | 0.143 |
| Map sharpening B factor (Å<sup>2</sup>) | -100 | -96 | -100 | -101 | -101 | -105 | -109 | -105 | -98 | -98 | -103 | -101 | -122 | -100 | -100 |

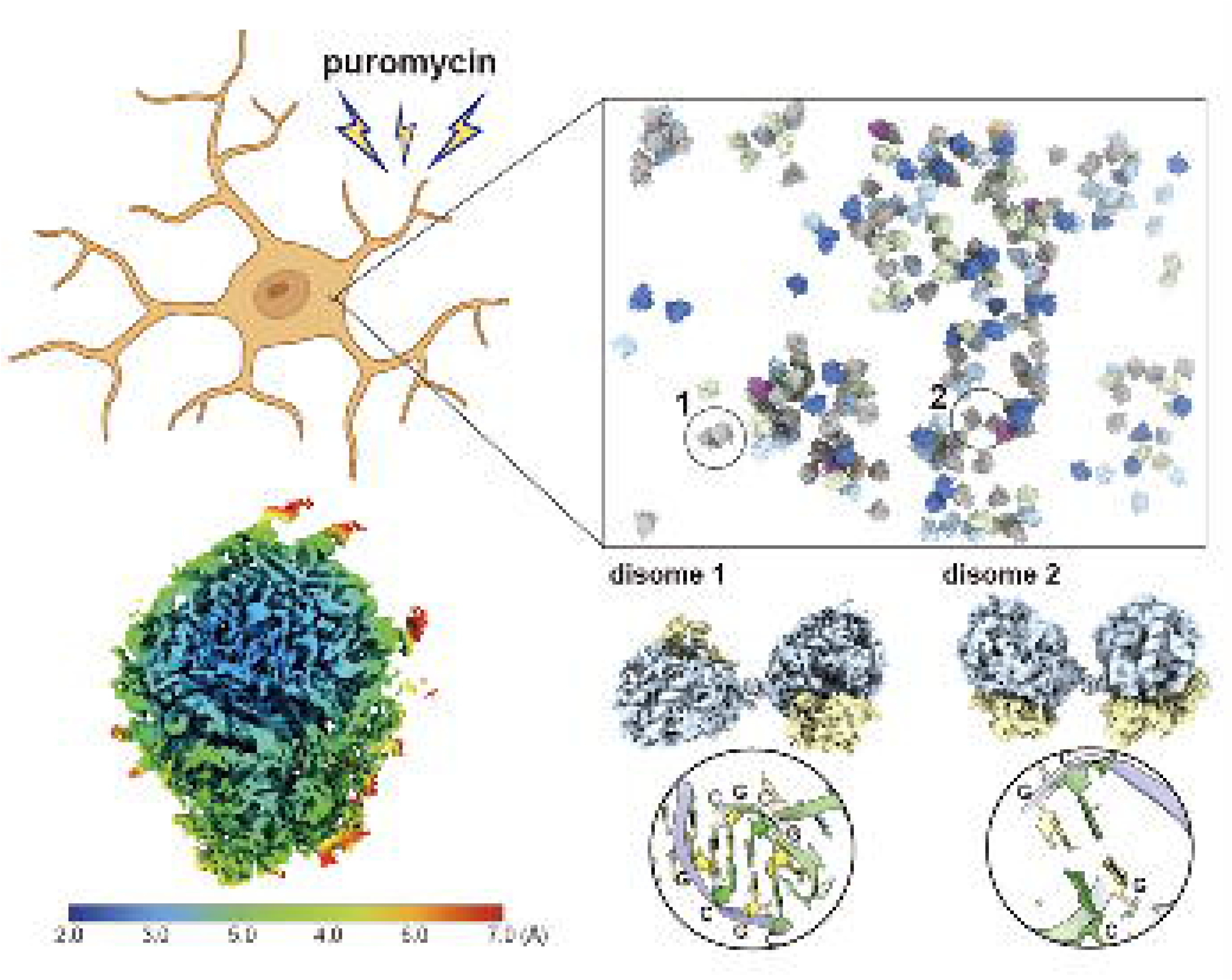

